# Internal gain modulations, but not changes in stimulus contrast, preserve the neural code

**DOI:** 10.1101/281444

**Authors:** Sangkyun Lee, Jiyoung Park, Stelios M. Smirnakis

## Abstract

Neurons in primary visual cortex (V1) are strongly modulated both by stimulus contrast and by fluctuations of internal inputs. An important question is whether population codes are preserved under these conditions. Changes in stimulus contrast are thought to leave population codes invariant, whereas the effect of internal gain modulations remains unknown. To address these questions we studied how the direction-of-motion of oriented gratings is encoded in layer 2/3 of mouse V1. Surprisingly, we found that, because contrast gain responses across cells are heterogeneous, a change in contrast alters the information distribution profile across cells leading to the failure of contrast invariance. Remarkably, internal input fluctuations that cause commensurate firing rate modulations at the single-cell level, do respect population code invariance. These observations have important implications for visual information encoding, and argue that the brain strives to maintain the stability of the neural code in the face of fluctuating internal inputs.

## Introduction

Contrast invariance of orientation or direction tuning functions, i.e., the preservation of tuning function shape across contrasts, is a fundamental property of visual neurons (Sclar G and RD Freeman 1982; Skottun BC et al. 1987). This suggests that the brain may use the same groups of cells to extract orientation or direction information across different visual contrasts. Accordingly, Busse et al. (2009) compared population responses across contrasts after averaging cell responses according to preferred orientation (Busse L et al. 2009) and concluded that the population code for orientation is preserved across contrasts. Later studies (Graf AB et al. 2011; Berens P et al. 2012) similarly found that neuronal pooling weights for orientation/direction decoding across contrasts are substantially preserved. A consensus has therefore been reached that the population code is preserved across contrasts. However, some other recent studies (Peirce JW 2007; Sani I et al. 2013) have reported that some cortical cells show larger responses to a range of intermediate contrasts than at 100% contrast; these intermediate-contrast selective cells may in theory encode more information at intermediate contrasts. These observations suggest that it is worth revisiting the concept of contrast invariance to ask specifically whether the population of cells that convey information about orientation or direction of motion remains identical across visual contrasts.

Neural responses are not modulated only by external stimuli. Internal inputs also modulate neural responses under identical external stimulation (Zohary E et al. 1994; Shadlen MN and WT Newsome 1998) changing neural population activity (Niell CM and MP Stryker 2010; Polack PO et al. 2013; Ecker AS et al. 2014; Reimer J et al. 2014; McGinley MJ, SV David, et al. 2015; McGinley MJ, M Vinck, et al. 2015; Vinck M et al. 2015). In fact, Fiser et al. have argued that most variability in the brain is due to internal activity, while sensory inputs evoke relatively small modulations superimposed on internally driven activity (Fiser J et al. 2004). Similarly, changes in behavior and brain state are known to modulate neuronal responses to identical stimuli (Niell CM and MP Stryker 2010; Polack PO *et al.* 2013; Ecker AS *et al.* 2014; Fu Y et al. 2014; Reimer J *et al.* 2014). These observations raise the question how the brain is able to maintain a stable representation of sensory information in the face of large internal fluctuations of neuronal activity. In particular, how internal fluctuations affect the population code for orientation or direction of motion remains an open question.

Below, we addressed these questions by studying the neural population code for moving oriented gratings in layer 2/3 of mouse area V1. We found that the performance of decoders remains essentially unchanged when they are trained and tested across different levels of spontaneously fluctuating internal input, whereas it degrades substantially when they are trained and tested across different stimulus contrasts. The substantial degradation of direction-of-motion decoders trained at different contrasts results primarily because the identity of cells that contribute most to direction decoding is not contrast invariant, but instead changes with contrast. We conclude that: 1) cortical circuits are optimized to maintain the stability of the neural code in the face of spontaneously fluctuating internal inputs, and 2) contrast invariance of the neural code fails substantially at the population level.

## Materials and Methods

### Animal preparation

All experiments and animal procedures were performed in accordance with guidelines of the National Institutes of Health for the care and use of laboratory animals and were approved by the IACUC at Baylor College of Medicine.

In our study, 9 C57/BL6 wild-type and 7 Thy1-GCaMP(Thy1-GCaMP6s 4.3(Dana H et al. 2014)) mice were used, which were 4-8 weeks old. During surgery, mice were anaesthetized with 1-1.5% isoflurane, and Baytril (5mg/kg), Carprofen (5mg/kg) and Dexamethasone (1.5mg/kg) were administered subcutaneously to minimize brain swelling (Holtmaat A et al. 2009). After attaching a headpost on the skull, a 3-mm diameter craniotomy was made on the center of visual cortex — 2.7mm lateral to the midline and 1.5mm posterior to the bregma. For the 9 wild-type mice, GCaMP6s virus (AAV5.Syn.Flex.GCaMP6s.WPRE.SV40, Penn Vector Core) was injected within the craniotomy by using a Drummond Nanojector (∼90 nl per site) after diluting 4-8 times with sterile saline. Then, the craniotomy was covered with a glass window.

### Imaging

Two-photon experiments were performed 3-4 weeks after the surgery, when GCamp6s expression is optimal. For Thy1-GCaMP mice, the experiments were conducted at 1-2 and/or 10 days after surgery without viral injection.

Populations of 50-200 cells located 150-250 µm below the pia were imaged with water-immersion objective lenses, either 20x, 0.95 NA (Olympus), or 16x, 0.8 NA (Nikon), in a modified Prairie Ultima IV two-photon laser scanning microscope (Bruker, Billerica, MA), fed by a Chameleon Ultra II laser (Coherent, Santa Clara, CA). Cell populations were imaged at frame rates of ∼7 Hz. Depending on imaging depth, the laser power was kept between 20 mW at the surface and 50 mW at depths below 200 µm, at 910 nm wavelength.

For experiments (n=19) with sedated animals (n=11), 1.5mg/kg of fentanyl and 0.5mg/kg dexmetetomidine were injected 0.5-1hr before the recording. Out of 28 main imaging sessions in different FOVs, 14 sessions (6 sessions from sedated animals) were performed with a time break (0.5-1 hours) between two sub-sessions with visual stimulation. During the time break, the screen was turned off and about 5-10 minutes before start of the second sub-session the screen was turned on again. During pre-/post-imaging sub-sessions, visual stimulation of 50-100 trials/condition was presented to mice. All the experiments were performed after verifying that neuronal population imaged responded to our visual stimulation though a brief retinotopy.

### Visual stimulation

Visual stimuli were generated in MATLAB and displayed using Psychtoolbox (Brainard DH 1997). Drifting at 2Hz, square-wave gratings at 0.04 cycles/degree were presented for 500 ms followed by an inter-stimulus interval of 1.5 seconds during which a full-field gray screen at the same mean luminance was presented. All trials (100-200 trials/condition) were pseudo-randomly interleaved. The stimuli were presented on an LCD monitor (Koolertron, Shenzhen, China) at 60 Hz frame rate, positioned 8 cm in front of the right eye, centered at 45 degrees clockwise from the mouse’s body axis. The visual angle of the screen spanned 56° elevation and 86° azimuth. The screen was gamma-corrected, and the mean luminance level used was 85 cd/m^2^. In some early experiments (n=8), another screen (DELL 2408WFP, Dell, Texas, USA) was used, of which visual angle spanned 54° (elevation) x 78°, at the mean luminance level of 80 cd/m^2^ after gamma correction.

For 28 imaging sessions gratings moving to 4 directions (−15° or −10°, 0°, 30°, and 90°) were presented randomly interleaved at 100%, 40% contrast, 100-200 trials per condition per session. For 12 sessions out of 28 sessions, 20% contrast was also used.

To assess contrast-dependent population codes for a small stimulus size, we performed 9 imaging sessions from additional 5 sedated animals. For these experiments, a small grating stimuli (i.e., 15 degrees in radius) moving to 3 directions (−30°, 30°, and 60°) were presented randomly interleaved at 100%, 30% contrast. The grating stimuli were presented on the aggregate receptive centers of cells imaged within an FOV.

Occasional experiments included grating stimuli spanning the full range of directions (0 to 330 degrees) at 30-degree intervals presented pseudo-randomly interleaved at 100% or 40% Michelson contrast (Michelson A 1927). Under these conditions, which were used to calculate full tuning functions, each stimulus was presented 30 times.

### Monitoring animal behavior

Animal behavior was monitored during awake experiments by tracking wheel rotation and recording ipsi-lateral eye movements (Suppl. Fig. 1). While the mouse head was restrained, the mouse was free to move forward or backward on the rotating wheel during experiments. The wheel rotations were measured with an incremental encoder with a resolution of 8000 cycles/revolution (Model 15T, www.encoder.com). Eye movements were monitored through a dichroic mirror (FM02, www.thorlabs.com), which was placed between the visual stimulation screen and the mouse eye, using an infrared camera (GC660, Allied Vision Technologies) at 30 frames/second (Suppl. Fig. 1A). Behavioral data acquisition was synchronized with the presentation of the visual stimulus and the acquisition of imaging frames.

### Data analysis

#### Preprocessing

Movies were x-y-motion-corrected by comparing image frames to the reference image with a sub-pixel registration method (Guizar-Sicairos M et al. 2008). For data from sedated animals, the average of the first 100 image frames was used as the reference image. For the awake data, 100 image frames for which no wheel movements appeared were used. For cell selection, a circular disk or annulus was manually defined to cover the viral expression over a cell body (Chen TW et al. 2013). After high-pass filtering to get rid of slow signal drifts (cutoff freq=0.05Hz), we corrected the neuropil contamination of the fluorescence signal (F) at the soma by subtracting the mean fluorescence of an adjacent neuropil patch annulus (extending from 7-20 ? away from the cell body center), F_n_, as follows: F_correct_ = F - S^*^F_n_ (Kerlin AM et al. 2010; Chen TW *et al.* 2013; Lee S et al. 2017), where S=0.65 similar to other studies (Chen TW *et al.* 2013; Dana H *et al.* 2014) using GCamP6 virus.

For awake experiments, sessions were first screened by the experimenter to exclude segments with obvious artifacts, such as eye squinting or inappropriate eye opening, excessive stress indicated by the restlessness of the animal. Data selected had to have successful monitoring of eye movements under good eye conditions (e.g., neither eye squinting nor inappropriate eye opening) for at least 30 minutes of visual stimulation. Then, a second pass of quantitative screening was performed. All trials with large movements were excluded from data analysis. Large movements were defined by either recording the instantaneous rotation speed of the wheel > 1cm/sec or the x-y movement of the image frame >2µm from the reference frame of each movie. When the movies in each experiment session show substantial z-drift, resulting in more than 10% cells identified in the first movie to disappear in the last movie, the entire session was excluded from analysis.

Eye position information was analyzed within the quiet awake state. Pupil size and location (x- and y-coordinate) were tracked with an automated custom-built program. Frame-by-frame supervised inspection of the eye traces followed by statistical analysis was then performed. In the following, statistical thresholds at each step were calculated within trials that survived in the prior steps. First, trials with eye movements whose velocity exceeded 2 standard deviations from the mean were excluded from analysis. Second, trials with large eye position deviations from the median were excluded from analysis. Specifically, trials were excluded when the X or Y coordinate was < 1st quartile coordinate - interquartile range (IQR), or > 3rd quartile coordinate + IQR. Typically, 90% of trials that survived this criterion showed less than 10.2° eye excursions from fixation (see example in Suppl. Fig. 1B; median excursion = 4.5° at this dataset), commensurate with ∼10% of the stimulus size (∼100° in diameter). This was consistent across sessions. Similarly, trials for which pupil size deviated more than the IQR away from the 1st or 3rd quartile were excluded from analysis, as they might correspond to relatively extreme states of arousal. Note that pupil size was not modulated as a function of stimulus contrast (Suppl. Fig. 1C-D), ensuring that contrast dependence of pupil size is not a confounding factor in our conclusions.

*Importantly, results obtained were consistent across awake and sedated animals, further ensuring that differences in eye movement profiles cannot explain our results.*

### Estimation of spike rates

To estimate the spike rate of each cell, the pre-processed fluorescence signal was normalized within that cell body, pixel by pixel, by calculating (F-F°)/F° (i.e., ΔF/F). For each pixel, F_0_ was defined as the mean fluorescence values over the time-series of that pixel. Spike rates were then estimated by applying a method (Lee S *et al.* 2017) we recently developed. Briefly, this method was based on a sparse non-negative linear regression model to estimate spike rates associated with the calcium fluorescence ΔF/F signal by assuming linear calcium dynamics with a time constant, which was adapted from (Chen TW *et al.* 2013), to represent the decay time of fluorescence signal in the cell body expressing GCaMP6s.

The linear relationship between fluorescence signal reflecting calcium dynamics and estimated spike was given as:

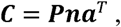

where ***C*** is a multi-pixel Matrix of [time-samples x pixels] of ΔF/F signals, ***P*** is a convolution matrix of [time-samples x time-samples] that generate the typical calcium dynamics from an spike-rate vector of [time-samples x 1], ***n***, and a is ***a*** spatial filter vector of [pixels x 1] that applies to across pixels used in the cell body. The superscript, *T*, represent the transpose of vector.

The convolution matrix was constructed by using the time constant (Chen TW *et al.* 2013) for GCaMP6s signal (i.e., τ=0.85 seconds) (see Lee et al.(Lee S *et al.* 2017) for more detail).

With constraining ***a*** ≥ ***o*** for the spatial filter and ***n*** ≥ ***o*** for spike rates, the objective function was given as:

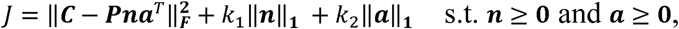

where *k*_1_ and *k*_2_ are the parameters that were automatically optimized while minimizing the objective function (see Lee et al.(Lee S *et al.* 2017) for more detail). We then find optimal a and n that minimize the cost function, where n yields the estimated spike rates for that cell.

### Measuring Visual Responses

For each cell, visual response was calculated trial-by-trial as the increase of the mean ΔF/F signal across pixels corresponding to the cell-body that occurs within the first 500ms of visual stimulation (compared to the 500ms immediately preceding the onset of the stimulus). The mean contrast-evoked responses of each cell were calculated by averaging all trial responses for that cell in each contrast, for 100% and 40% contrasts, respectively. To be included in subsequent analysis, the contrast-evoked response of a cell had to be greater than 5% at either contrast. In the results we present, ‘all cells’ refers to all the visually responsive cells that pass this criterion.

For selected cells, single trial responses were calculated for each trial as the average spike response from a 4-frame window closely matching the visual stimulus duration. For awake animals, the window computed for the trial response was the same as the window for visual stimulation. For sedated animals, it was centered to the peak of the average visual response (see Fig. 1D), as the time course of visually evoked responses were somewhat prolonged in sedated animals (Haider B et al. 2013).

**Figure 1:**
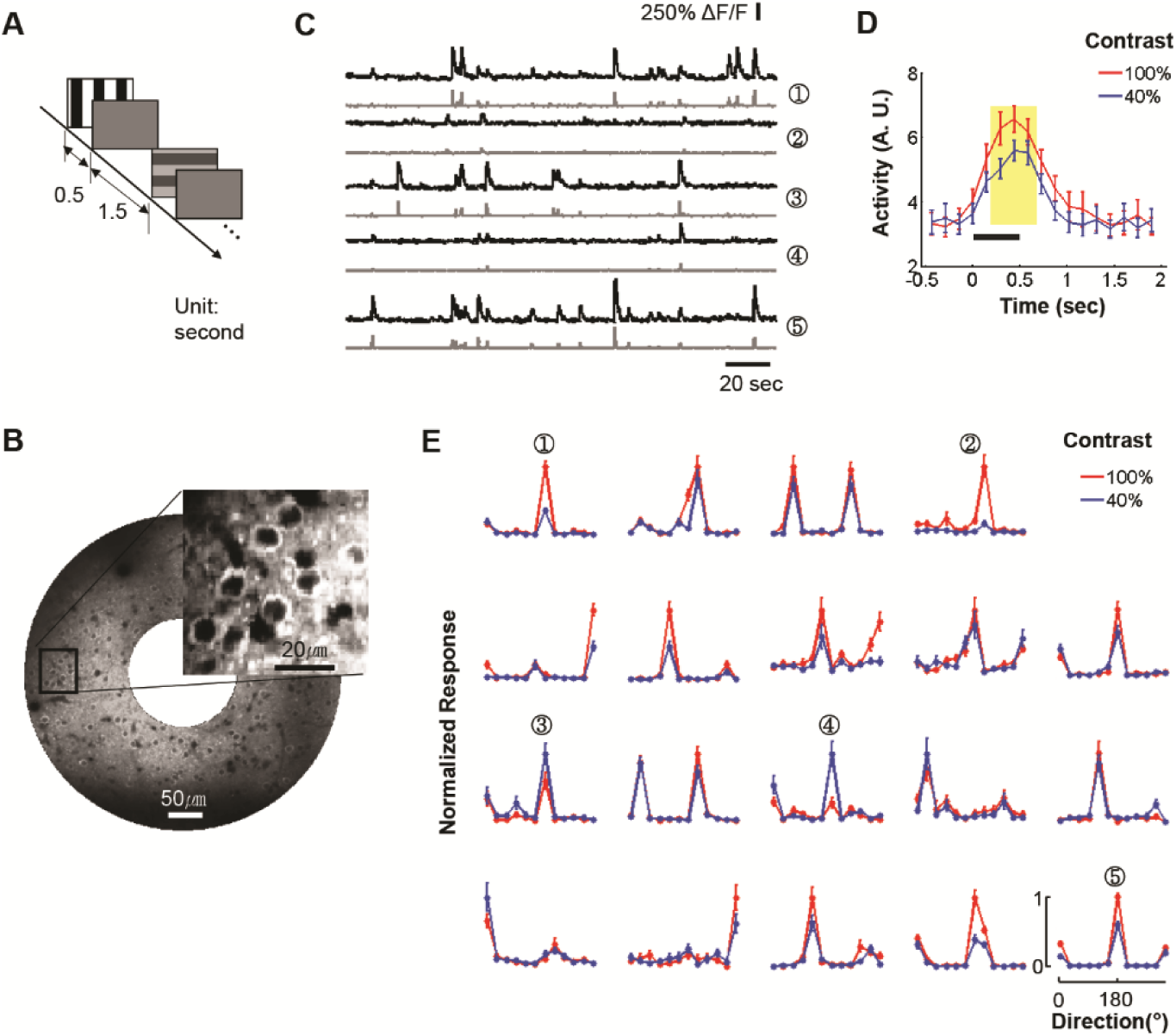
Direction tuning functions across cells scale differently with contrast. (A) Stimulation paradigm: In each trial, a grating drifting in one of twelve directions was presented for 500 milliseconds followed with 1.5 sec of uniform illumination at the same mean luminance. Direction of motion and contrast conditions were randomly interleaved across trials. **(B)** Mean fluorescence image from a Field-of-View that expresses GCaMP6s. Inset: enlarged view from the indicated rectangle. **(C)** Examples of fluorescence traces (top) and corresponding deconvolved spike train activity (bottom). **(D)** Mean visual spike responses across all stimulus directions (n=12) and all cells (n=102) analyzed in this FOV. Black bar: period of visual stimulation. Yellow shade: frames used for the calculation of trial responses. Mean±SEM. **(E)** Direction tuning curves normalized by the maximum response at 100% contrast for each cell separately. Cells 1-5 correspond to the traces shown in **(C)**. Mean±SEM (n=30 trials/direction). Note that while the preferred direction of cells is well preserved across contrasts, the relative scale of the response (gain) varies widely across cells. See for example cells 3, 4 whose responses to lower contrast are higher than to 100% contrast. Note that this is not the result of poor signal to noise ratio (as shown by the good quality of the recording traces in **C**).

### Decoding stimulus direction from population activity

To discriminate the visual stimulus direction from cell population responses within versus across contrasts, we used a regularized logistic regression model (RLRM) with the L2-norm regularization (Krishnapuram B et al. 2005; Bishop CM 2006). In this model, the cells’ trial-response vector was fed into an input vector 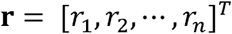 of RLRM, where r_i_ is the *i*-th cell trial-response in the given trial.

Then, the input vector was classified with the RLRM model as follows:

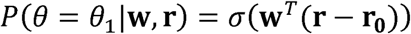

Here, *σ* (*x*) = 1/(1 + exp(-*x*)), 0 is stimulus direction, **w** is a weight vector, and

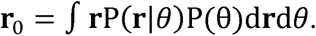

For the binary classification with equal number of trials per condition,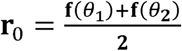, where

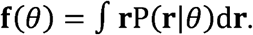

We performed a mean response subtraction (i.e., **r** – **r**_0_) to achieve an unbiased decoder (i.e.,∫**w**^T^(**r** – **r**_°_)P(**r**|*θ*)P(θ)d**r**d*θ*=0). Our unbiased decoder is comparable to Graf et al. (Graf AB *et al.* 2011) that applied contrast-response-offset correction. Note also that our conclusions shown in the present study remained the same without this bias correction and for z-normalization of individual cell responses.

The L2-norm regularization on w was applied to achieve a maximum a posteriori (MAP) estimate as follows:

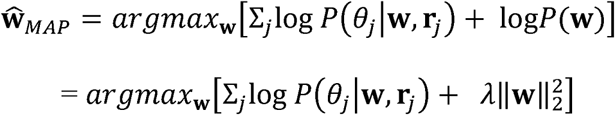

Here, λ is a free parameter searched in a logarithmically linear space between 10^−10^ and 10 to find the optimal value that results in the maximum decoding performance, using a separate set of data from the training and testing data (see below).

The decoding performance of cell populations for a pair of stimulus directions were assessed and cross-validated by 100 random sub-sampling tests (80% for training, 10% for optimal λ searching, 10% for testing). The decoding tests were performed: 1) within the same contrast as the training contrast to assess decoding performance for within-contrast direction decoders, or 2) within a contrast different from the training contrast for cross-contrast direction decoders. The same method was also applied to compare decoding performance within versus between population-activity-levels (PALs). To this end, trials within each stimulus direction and within each contrast were separated into two sub-groups, one with high (50%-highest, “**H**”), the other with low (50%-lowest, “**L**”) average population activity. The average population activity for each trial was computed as the mean response across all cells included within each FOV.

The logistic regression model (LRM) was also used for selecting the most informative cells in stimulus-direction decoding. This was achieved by using an L1-norm regularization technique (instead of the L2-norm) to increase the sparseness of the weight vector (sparse LRM). We then re-evaluated the decoding performance with L2-norm RLRMs through cross-validation for the selected cells. Specifically, we trained a sparse LRM from 80%, randomly selected, training data trials using 10% distinct data trials for λ optimization. A number (n) of cells with the n-highest magnitude of weights were then selected. The contribution of each cell to the output value of LRM before the nonlinear function (i.e., **w**^*T*^(**r** – **r**_0_)) was determined by two factors: 1) the overall response modulation of the cell between conditions used, and 2) its corresponding weight value, *w*_i_. The different response modulations across cells were normalized by setting the response variance for each cell across trials to unity prior to training in order to fully reflect the extent of cell’s contribution to the decoder. Following cell selection, an L2-norm regularized RLRM was trained to find the optimal λ and re-optimize the weights of the selected n-cells using new randomly selected training data (80% trial), as well as distinct λ optimization (10% of trials), and distinct testing (10% of trials) data. This process was repeated 100 times. Note that the data for cell selection and decoder training with an optimal λ were kept strictly separate from the testing data.

### Linear Direction Tuning Function Fits across contrasts or different population activity levels

To obtain the response gain of direction tuning functions of each cell between 100% and 40% contrast, we used a linear fit as follows:

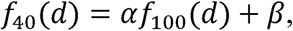

where *f*_4O_(d) and *f*_100_(d) represent the tuning functions of a cell at 40% and 100% contrast respectively, *d* refers a stimulus direction, and *α* and *β* represent a gain and a bias.

To prevent an arbitrary non-physiological fitting (e.g., a negative *α* or a large positive *α* with a large negative *β*), we constrained the fitting as follows:

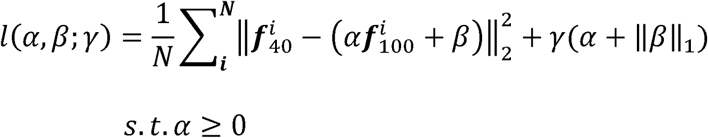

To minimize the cost function with the non-negative constraints, we adapted a log-barrier technique (Boyd SP and L Vandenberghe 2004; Kim S-J et al. 2007).

Here,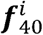 and 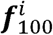 represent a vector composed of the mean evoked responses, 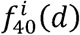 and 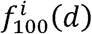, from the *i*^th^ sub-sampled set of trials of each cell to stimulus direction (*d*) at 40% and 100% contrast, respectively, and γ is a free parameter that constrains the magnitude of *α*and *β*. In the *i*^th^ sub-sampling, 50% trials randomly selected within stimulus direction=*d* and contrast=*c* were averaged to generate 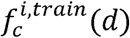and the remaining 50% trials to generate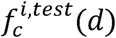(d). This random subsampling was performed N=1000 times, i.e., *i*=1,…,N. Then, the training set composed of 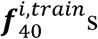 and 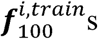 was used to search for the optimal γ in a logarithmically linear space between 10^−5^ and 10 and to estimate the parameters *α* and *β*. The testing set of 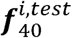and 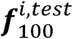 was used to evaluate the fit’s performance. The same strategy was also applied to obtain the response gains when comparing between different population-activity-levels by conditioning trials based on population-activity-level as well as contrast (e.g., ***f* _*100L*_** = *α****f* _*100H*_** + *β*).

For within-contrast fits (e.g., ***f* _*100*_** = *α****f* _*100*_** + *β*), training trials obtained from the *i*^th^ sub-sampling shown above were further divided into two subsets to construct Y, X (i.e., Y = aX+ /3) by averaging 50% trials randomly selected from training trials within each stimulus condition for Y and the remaining 50% trials for X. The same method was applied for testing data.

The bias *β* was normalized to the maximum response at 100% contrast before plotting in Fig. 2B-C and 5E.

**Figure 2:**
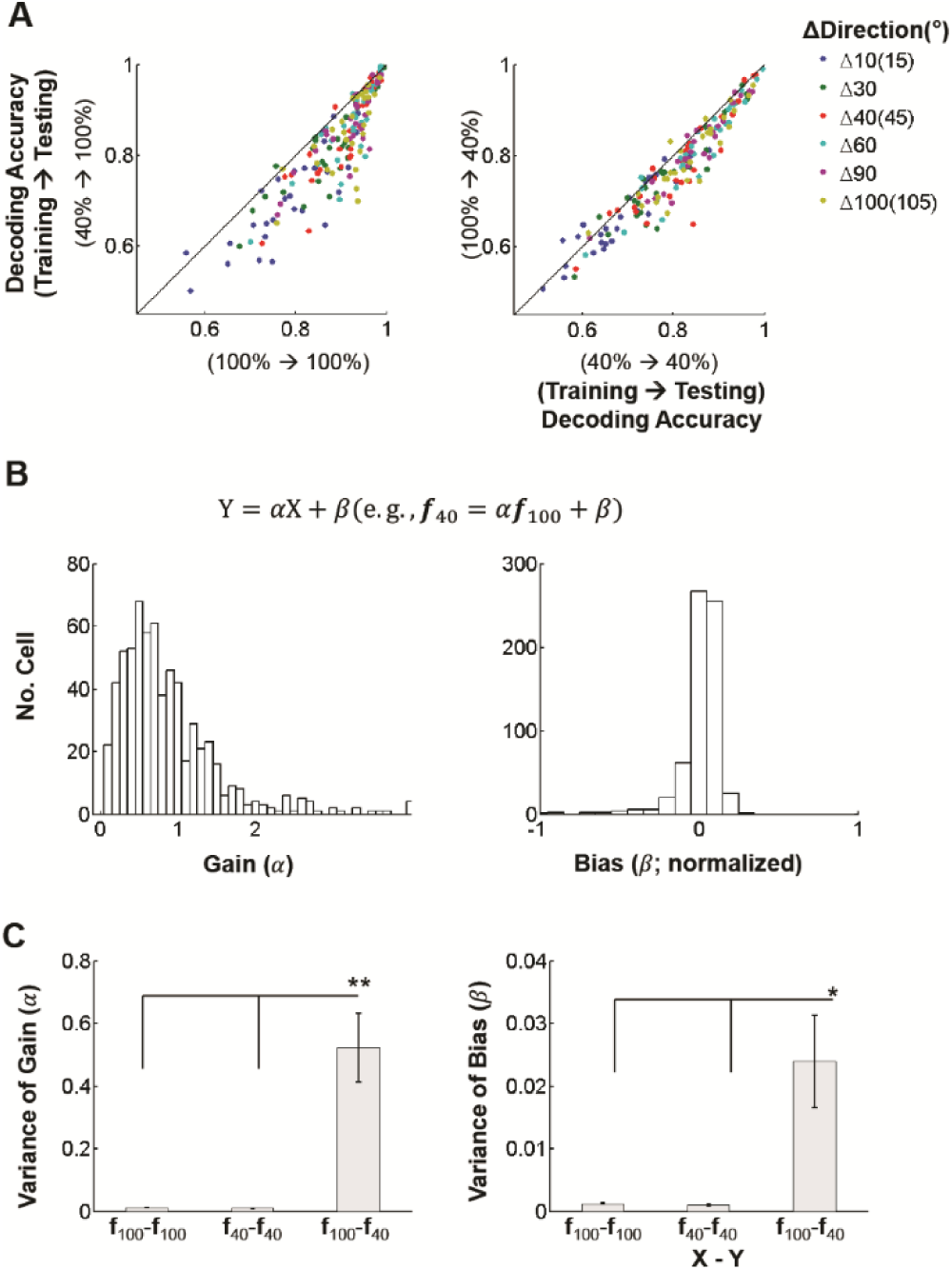
Direction population code violates contrast invariance. (**A**) Within-contrast decoders (X-axis) versus cross-contrast (Y-axis) decoders tested at 100% (*left panel*; P<1e-9, Friedman Test) and 40% contrast (*right panel*; P<2e-3). Each dot represents decoding accuracy from a single FOV (n=28). Colors represent the difference between decoded directions of stimulus motion (in degrees). (**B**) Distribution of gain (*α*) and bias (*β*) across FOVs (n=28) in the linear fit of ***f***_40_ = *α****f***_100_ + *β*: *α* (*left panel*) and *β* (*right panel*). *β* was normalized to the maximum tuning response at 100% contrast. Large dispersion of contrast gain parameters occurs across cells. (**C**) The mean variance of the extracted parameters across FOVs (n=27) when fitted within (**f**_100_-**f**_100_, **f**_40_-**f**_40_) versus across contrasts (**f**_100_-**f**_40_). Only cells whose fits had explained variance >0.5 were included (one FOV whose cells showed lower explained variance than the threshold was excluded). /3 was normalized to the maximum tuning response at 100% contrast. These plots show that the large dispersion of parameters across cells for the **f**_100_-**f**_40_ contrast transitions represents a physiological effect and does not arise as a result of variability of sampling. Error bar: SEM. P<1e-10, Kruskall-Wallis Test for *α* (*left panel*) and *β* (*right panel*). ^*^,^**^: P<0.01, P<5e-5 in post-hoc Tukey tests.

### Identifying which Cells Contribute to Direction Decoding Across Conditions

*P(c*_*i*_*|contrast, n)* was defined as the probability that the cell *c*_*i*_ belongs to the *n* “most informative” cells, i.e. the cells whose output can discriminate best between the stimulus conditions at the given *contrast*. Practically, *P(c*_*i*_*|contrast, n)* was calculated by counting how many times out of 100 cross-validation tests, the cell c_i_ was selected as one of the *n* “most informative” cells. Then, the probability for the cell to belong to the *n* most informative cells at both 100% and 40% contrast was given as: 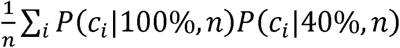. We considered cells to be reliably selected among the *n* “most informative” cells, if either *P(c*_*i*_|100%,*n*) or *(c*_*i*_*|40%,n) > 0.7.* The probability for a cell to reliably belong to the *n* “most informative” cells at both contrasts is then:

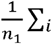sign(P(c_i_|100%, n))sign(P(c_i_|40%, n)), where n_1_ = minimum(n_100_, n_40_), and n_100_, n_40_ represent the number of cells with P(c_i_|contrast, n) > 0.7 at contrasts 100% and 40% respectively.

### Decoding Performance in Control Data Constructed to have identical SNR across different contrasts

To rule out the possibility that contrast-dependent difference in SNR affects the comparison between within-contrast versus cross-contrast decoders, we generated artificial control data by modifying relative noise levels (i.e., the SNR) of 100%-contrast data to be the same as the one of 40%-contrast data within each stimulus direction for each cell. At the same time we held the mean response within stimulus conditions the same in order to maintain original direction tuning functions and contrast response gains. We then compared the decoding performance of within-contrast versus cross-contrast decoders derived from these control data.

To generate spike-rate responses, we employed the Gamma distribution. We chose the Gamma distribution rather than Poisson distribution because spike response variance on repeated identical stimulus presentation is often larger than predicted by the Poisson distribution, whose relative dispersion is constant (i.e., variance/mean = 1) This is supported by a recent study, which found the negative binomial distribution (NBD) to be a better model of real spike fluctuations (Goris RL et al. 2014) (the Gamma distribution resembles the behavior of NBD for estimated relative spike-rates). The mean and variance of the response of each cell within each stimulus condition was calculated at 100% and 40% contrast, respectively. These values from each contrast were then used to create a gamma distribution for each cell, contrast as follows:

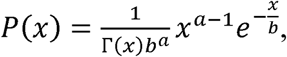

where 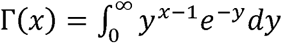, where x represents the firing rate. The two free parameters, *a* and *b*, determine the mean and the variance as *ab* and *ab*^*2*^ respectively. Using this relationship, we generated 1000 random samples for each contrast, with a and b given by *a*_*40*_and 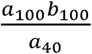for 100% contrast and *a*_*40*_ and *b*_*40*_ for 40% contrast, respectively. The subscript indicates the contrast used to estimate the corresponding parameter. For 40% contrast, this process generates artificial data that have the same mean and variance as the original data, and for 100% contrast it generates data that have the same original mean obtained at 100% contrast, but signal to noise ratio that matched the signal to noise ratio obtained at 40% contrast. This was performed cell-by-cell across all cells. The entire process was performed 10 times to generate data for cross-validation. The usual procedure was then applied to evaluate decoder performance within- and across contrasts, as described above.

### Statistical Analysis

All statistical tests were performed with non-parametric, MATLAB built-in functions. Post-hoc Tukey test refers the ‘tukey-kramer’ method in the function called ‘multcompare’. FWE and SEM in figure legends represent the Bonferroni family-wise-error correction and the standard error of the mean.

## Results

### Contrast gain responses are heterogeneous

We imaged neurons in layer 2/3 of mouse V1 via 2-photon microscopy (Fig. 1B) while presenting oriented gratings drifting in one of 12 different directions, at 100% or 40% contrast. Direction tuning functions were calculated per contrast condition after deconvolving the calcium fluorescence (ΔF/F) signal (Figs. 1A-D). Tuning function shape is preserved across contrasts (Fig. 1E), as shown previously in cats (Hubel DH and TN Wiesel 1959; Sclar G and RD Freeman 1982) and monkeys (Hubel DH and TN Wiesel 1968). However, response gains vary considerably across cells as a function of contrast (Fig. 1E; Suppl. Fig. 2). Some neurons even exhibit larger preferred orientation responses at 40% versus 100% contrast (Fig. 1E; cells #3, 4). The heterogeneity of contrast gain responses across cells raises the possibility that different optimal population codes for direction-of-motion are implemented at different contrasts.

### Population codes for direction depend on visual contrast

To test whether population codes for direction-of-motion are contrast-invariant, we collected 100-200 trials per stimulus condition for stimuli spanning 4 directions. Twenty eight imaging sessions from different fields-of-view (FOVs) were performed in 11 sedated and 6 awake animals (50-150 neurons/FOV; see Methods). All sessions (18, 10 FOVs from sedated, awake animals respectively) were grouped together, as sedated and awake animals gave similar results. A fraction of the trials was used to train a decoder to classify stimulus direction from population responses trial-by-trial. The mean decoding accuracy was then obtained from 100 cross-validation tests, in which all training and testing data were exclusively separated. Decoding accuracy across FOV’s ranged from 88% (100%-contrast) to 80% (40%-contrast), confirming data quality (Suppl. Fig. 3A).

To examine whether population codes are preserved across contrasts, we compared how the decoder performed when training and testing trial sets were taken from the same versus across different contrasts. Contrast invariance predicts that decoding performance should be independent of training contrast. In contrast, we found that decoding accuracy was better within than across contrasts (Fig. 2A and Suppl. Fig. 3B). Specifically, when testing with 100%-contrast data, training the decoder with 100%-contrast data outperformed training with 40%-contrast data (Fig. 2A *left*; P<1e-9). Conversely, when testing with 40%-contrast data, training with 40%-contrast data outperformed training with 100%-contrast data (Fig. 2A right; P<2e-3). This was not due to data overfitting or differences in signal to noise ratio, as the decoders obtained remained robust across acquisition time shifts in the training/testing data sets (Suppl. Fig. 3C) confirming the robustness of our decoding strategy. The fact that within-contrast decoders outperform cross-contrast decoders is a signature of the limits of invariance of the population direction code across contrasts, and suggests that contrast-specific codes might be the underlying rule for optimal population coding.

### Different cell ensembles contribute to direction encoding at different contrasts

The degradation of cross-contrast direction decoding performance has its roots in the heterogeneity of contrast gain responses across neurons (Fig. 2B). We fit the direction tuning function, ***f***_40_, at 40% contrast to the one, ***f***_100_, at 100% contrast as: ***f***_40_ = *α****f*_100_** + *β* (methods).The gain parameter *(α)* shows large variability (0-2) across cells, even when fits have high explained variance (> 0.5; Fig. 2B *left*), whereas the bias parameter *(α)* remains concentrated near 0 (Fig. 2B *right*). This was consistent with the fits from full tuning curves (Suppl. Fig. 2). The much tighter dispersion of both the parameters seen within contrast persists when fitting data from distinct time periods but the same contrast (Fig. 2C), as well as when applying a more conservative cell selection (Suppl. Fig. 3D). The marked heterogeneity of response gains and, to a lesser degree, biases across contrasts argues that different cells contribute differently to direction-of-motion encoding across contrasts, suggesting a change in the population code.

In order to test this directly, we used a sparse logistic regression model (LRM) to decode the stimulus direction of motion from the neuronal population activity in layer 2/3 of area V1. Cells were ranked from most to least “informative” depending on the magnitude of the weight with which they contributed (see Methods). There was only 25% chance that the same cell would be selected as “most informative” at both 100% and 40% contrast (Fig. 3A and Suppl. Fig. 4). Fig.3A plots the probability that a cell is selected as one of the ***n*** “most informative” cells for both contrasts as a function of ***n***. Although this probability naturally increases with ***n***, it still remains <50% at ***n***=20, although decoding performance is nearly plateaued (Fig. 3D). Figs. 3B-C show an example of contributing cells and their corresponding tuning functions for decoding gratings moving at 0° versus 30°, when ***n***=3. This example shows that cell 2, which was almost exclusively selected at 40% contrast (Fig. 3B), responds stronger to 40% contrast than to 100% contrast at direction=30° (Fig. 3C). These observations suggest that direction-of-motion information across contrasts is carried by substantially different populations of cells.

We further compared the amount of information encoded by the most informative cells for within-contrast (WC) versus cross-contrast (CC) direction decoding. Briefly, from the most informative ‘***n***’ cells at a given contrast, a regularized LRM was trained to re-optimize the weights only within the cells selected, and then used to test decoding performance within versus across contrast (see Methods; distinct data were used for testing, cell-selection, and classifier-training). As expected, decoding performance increases with number of cells (***n***) but, consistent with Fig. 2, remains significantly better within versus across contrasts for all ***n*** (Fig. 3D).

**Figure 3:**
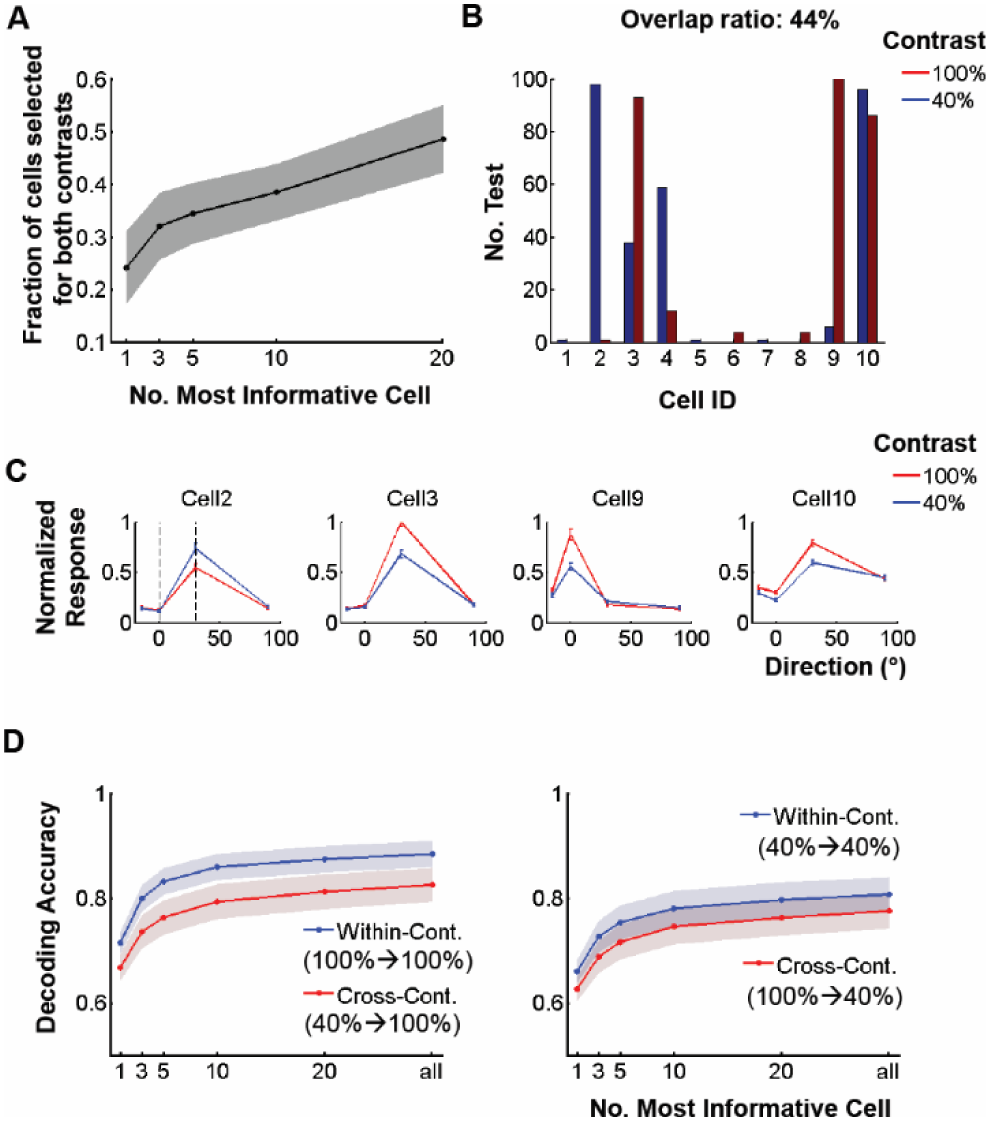
Cells that participate in direction decoding at different contrasts are substantially different. (**A**) Probability for a cell to belong to the first n “most informative” cells both at 100% and at 40% contrast. (**B**) *<u>Example:</u>* number out of 100 cross-validation tests for which cells belong to the first 3 “most informative” cells for decoding a change of direction =30°. (**C**) Direction tuning functions of cells selected in (**B**), normalized to the maximum response of cell #3. Mean± SEM. The two vertical dash lines represent the two directions decoded. (**D**) Decoding accuracy within (100%->100%; 40%->40%) versus across (40%->100%; 100%->40%) contrasts using only the first “*n*” most informative cells as a function of n. Friedman Test P<1e-10 (*left panel*), P<1e-5 (*right panel*). For (**A**) and (**D**), solid lines and shadows represent mean (n=28) and 95% confidence intervals, respectively.

### Contrast-dependent differences in noise characteristics do not explain away contrast-specific codes

To ensure that different noise levels at 40% vs 100% contrast are not the source of the decoding differences observed, we modified the data cell by cell, keeping the original mean responses, but adjusting noise levels so that SNR remains invariant across contrasts. We again found that within-contrast decoders outperform cross-contrast decoders, reinforcing the conclusion that this effect (Fig. 4A) is not due to different SNR levels across contrasts, but rather to the heterogeneity of contrast gain responses across the population of cells (Figs. 2B-C). Neither does destroying noise-correlation structure by randomly shuffling trials, cell by cell, within each stimulus condition influence our result (Fig. 4B).

**Figure 4:**
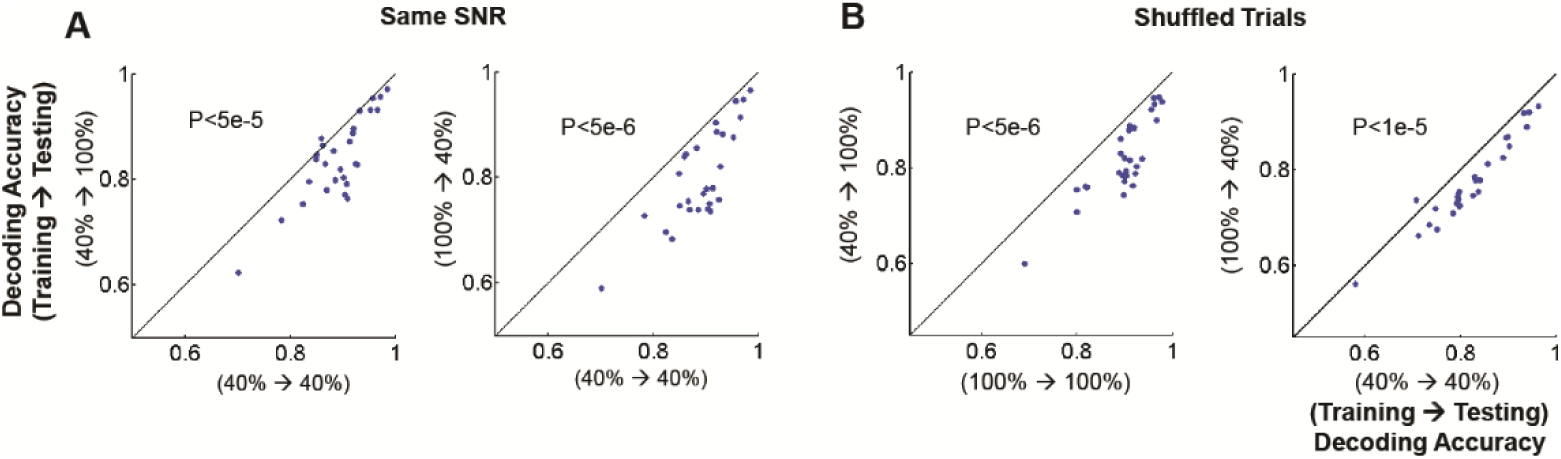
Better performance of within-contrast decoders does not result from contrast-dependent noise characteristics. Decoding performance of within-contrast (X-axis) versus cross-contrast (Y-axis) decoders after adjusting the signal to noise ratio of 100%-contrast data to match that of 40%-contrast data. *Left panel (Y-axis):* 100% contrast data are used for testing, 40% for training. *Right panel (Y-axis):* 40% for testing, 100% for training. (**B**) Decoding performance of within-contrast (X-axis) versus cross-contrast (Y-axis) decoders after we destroy the noise correlation structure across neurons. *Left panel:* testing contrast 100%. *Right panel:* Testing contrast 40%. To destroy noise correlations trials were shuffled within stimulus conditions and within cells (see Methods). Each dot indicates average decoding accuracy across all pairs of stimulus directions. Statistical test: Wilcoxon signed-rank test.

### Contrast-specific decoders still outperform cross-contrast ones for a small stimulus

Neuronal response decreases as stimulus size increases beyond their receptive field size (Allman J et al. 1985). This effect is more obvious for high contrast (Sceniak MP et al. 1999) and thus intermediate contrast stimulation may evoke stronger responses than high contrast stimulation in some cells. Therefore, one may wonder whether our finding is only valid when the large stimulus size was used. To this end, we stimulated cells by presenting on the aggregate center of their receptive fields a small stimulus (i.e., 15 degrees in radius), which was only slightly larger than the typical receptive field size (i.e., ∼10-12 degrees; Smith and Hausser (Smith SL and M Hausser 2010)) of mouse V1 cells. We also lowered the low contrast condition to 30% to increase the number of cells that do not exhibit surround suppression. We found that contrast-specific decoders outperformed cross-contrast ones (Suppl. Fig. 5), as they had with the full field stimulus. Therefore contrast-dependent surround suppression effects do not explain away out observations.

### Contrast-specific decoders outperform contrast-independent decoders

Finally, to test whether a universal pooling rule (decoder) could perform well across contrasts, we trained classifiers using data from all contrasts together then tested them at one of the contrasts. Our analysis again showed that contrast-specific decoders outperform contrast-independent decoders (Suppl. Fig. 6).

These observations strongly suggest that the population code for the direction-of-motion of moving gratings is not strictly invariant across modulations of stimulus contrast. We argue that this is due to the heterogeneity of contrast gain responses across cells (Fig. 2B, C). An important question is whether the code is better preserved during internal input fluctuations, which often result in similar or even greater modulations of firing rate and are ubiquitous across the brain (Fiser J *et al.* 2004).

### Spontaneous internal modulations leave population codes invariant

To probe whether the population code for direction-of-motion changes with internal input modulations that occur spontaneously, we separated the data trial-by-trial based on aggregate population activity levels (PALs). Within each stimulus condition trials were separated into two sub-categories: One with high (50%-highest, “**H**”), the other with low (50%-lowest, “**L**”) average population activity (see Methods). Surprisingly, decoding accuracies were essentially identical when the decoder was trained/tested within versus across these sub-categories. This was true for trials at both 100% and 40% contrast (Figs. 5A-D). Significant changes in decoding accuracy were only seen across changes of stimulus contrast, not when aggregate population response changed due to the spontaneous fluctuation of internal inputs (Figs. 5A-D). This was further corroborated by comparing the pattern of cells that contributed significantly to direction decoding across i) changes of stimulus contrast, and ii) different population activity levels (PALs). As expected, the pattern of cells contributing significantly to decoding was more similar between high and low PALs than between 100% and 40% stimulus contrast (Suppl. Fig. 7). The relative preservation of population codes across spontaneous fluctuations of population activity is closely related to the fact that response gain modulations across V1 L2/3 cells are considerably more uniform under this condition (Fig. 5E).

Note that, individual cell firing rate fluctuations from trial to trial within each stimulus condition are more extreme during spontaneous internal input modulations than across changes in stimulus contrast (Fig. 5F). In descending order population response activity levels were: (100%-contrast;**H**) > (40%-contrast; **H**) >> (100%-contrast; **L**) > (40%-contrast; **L)**. Therefore the differences noted cannot be attributed to potentially weaker modulations occurring during the spontaneous fluctuation condition. In addition, the mean signal-to-noise ratio across cells was preserved regardless of global internal modulation within each contrast (Fig. 5G). Therefore these factors do not alter our basic conclusions.

**Figure 5:**
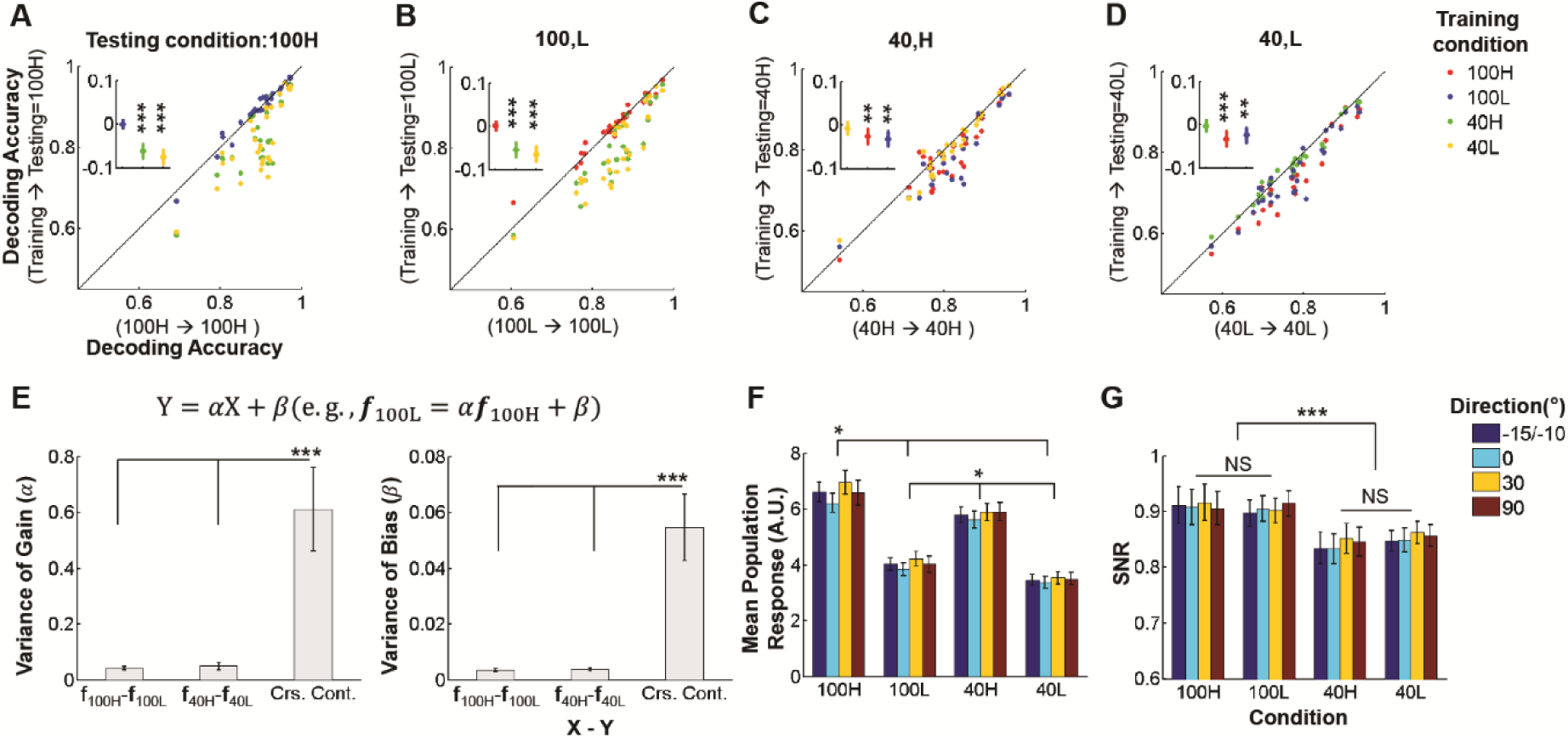
Internal gain modulations preserve the population code for stimulus direction-of-motion. (**A-D**) Decoding accuracy comparisons when training/testing data are taken across different stimulus contrasts and population-activity-levels (PALs). *<u>Inset:</u>* Cross-condition decoding accuracy minus within-condition decoding accuracy. ^**^: P<1e-3, ^***^: P<1e-4 (FWE, Wilcoxon signed-rank test). Note that population code is preserved across *H*igh ↔ *Low* internal gain modulations occurring at fixed stimulus contrast, but not across different contrasts. (**E**) Variance of gain (*left panel*) and bias (*right panel*) modulations across L2/3 cells, arising as a result of a change in population activity level (i.e., **f**_100H_-**f**_100L_, **f**_40H_-**f**_40L_) versus stimulus contrast (Crs. Cont.). ‘Crs. Cont.’ refers to the average variance derived from cross-contrast-fits (i.e., **f**_100H_-**f**_40H_, **f**_100H_-**f**_40L_, **f**_100L_-**f**_40H_, **f**_100L_-**f**_40L_). Cells with explained variance >0.5 were included in the calculation of variance. One FOV was excluded in the analysis due to low explained variance. P<1e-10, Kruskall-Wallis Test for *α* and *β*. ^***^: P<1e-7 in post-hoc Tukey tests. Note that *H*igh ↔ *Low* internal activity fluctuations lead to more homogeneous gain modulations across the L2/3 cell population than changes in stimulus contrast. (**F**) Mean population response amplitudes (arbitraty units) as a function of contrast (100% vs 40%) and population response level (High vs Low). P<5e-10 in Kruskall-Wallis Test after averaging across stimulus directions. ^*^: P<1e-2 in post-hoc Tukey tests. (**G**) Corresponding Signal-to-Noise Ratios (i.e., mean/standard deviation). Signal to noise ratios do not vary across internal modulation states, within a given stimulus contrast: P=0.84, 0.31 for 100H vs 100L, 40H vs 40L respectively. ^***^: P<5e-6 across stimulus contrasts: i.e., mean(100H,100L) vs mean(40H,40L); Wilcoxon signed-rank test after averaging across stimulus directions. NS: Not Significant. Mean ± SEM (n=28) for all error-bar plots except **E** (n =27).

In brief, spontaneous internal input fluctuations result in more homogeneous gain fluctuations across L2/3 V1 cells thereby preserving the population code invariance for direction-of-motion.

We further compared decoding performance by the *n* most informative cells as a function of *n*, similar to Fig. 3D. Decoding performance by the cross-condition decoder was degraded more strongly by modulations of stimulus contrast compared to spontaneous modulations of internal input (Figs. 6A-D). Interestingly, cells selected as 10 most informative at 40% contrast show on average ∼22% higher contrast gain at 40% contrast compared to cells that were selected as most informative at 100% contrast (Suppl. Fig. 8 and see also the accompanying text in the supplementary material). This again confirms that significantly different populations of cells contribute to direction-decoding at the two different contrast levels. In contrast, cells selected as most informative at low population activity levels (PALs) showed on average only ∼5-6% higher gain compared to cells selected as most informative at high PALs within each contrast condition.

**Figure 6:**
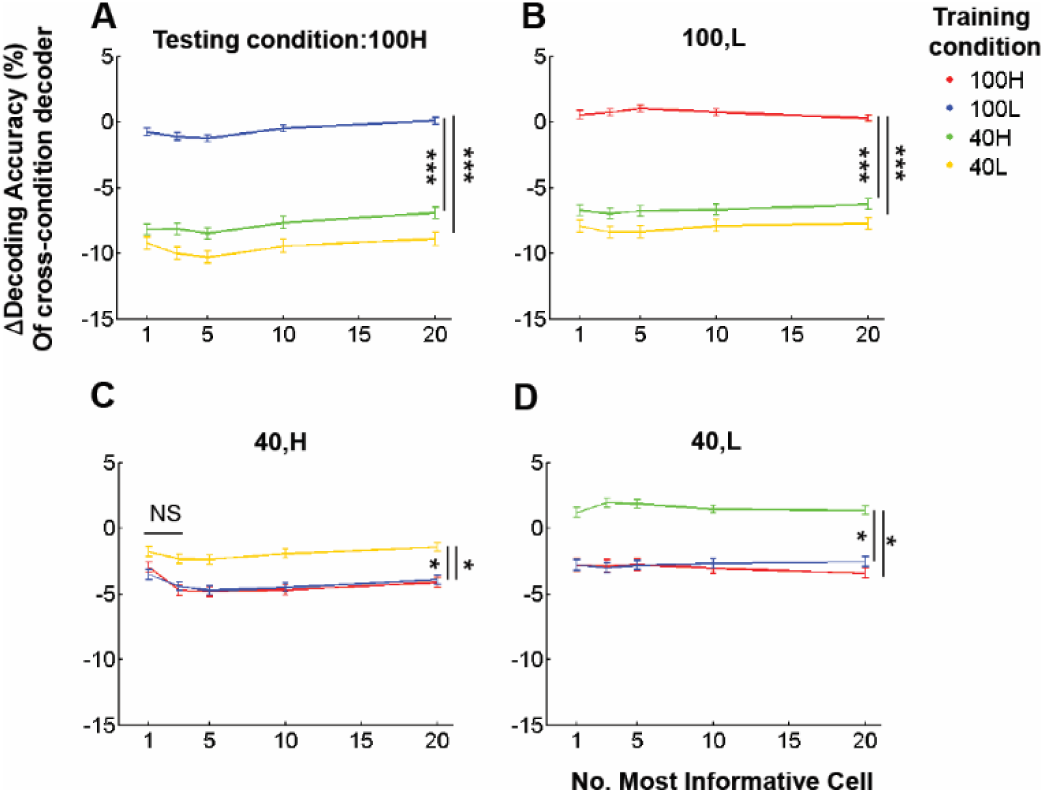
Decoding performance degrades in changes in contrast, but not in population activity. (**A-D**): Relative performance of cross-condition decoders generated using the first “n” most informative cells, as a function of “n.” Y-axis plots change in decoding accuracy: (cross-condition – within-condition)/within-condition. Illustrated differences are significant for all n, except for n=1,3 in panel (c): Kruskall-Wallis Test P<1e-7 (**A-B**), P<0.05 (**C-D**). ^*^P<0.05, ^***^P<1e-5 by post-hoc Tukey tests.

## Discussion

### Changes in Stimulus Contrast do not Leave Population Codes Invariant

We chose to compare 100% to 40% contrast because mouse V1 neurons respond well and mice discriminate orientations successfully (Long M et al. 2015) at these contrasts. The functional properties of mouse V1 neurons observed in the present study agree with earlier studies (Albrecht DG and DB Hamilton 1982; Ohzawa I et al. 1982; Sclar G and RD Freeman 1982; Ohzawa I et al. 1985; Skottun BC *et al.* 1987; Geisler WS and DG Albrecht 1992). First, the shape of direction tuning functions was contrast invariant (Sclar G and RD Freeman 1982; Skottun BC *et al.* 1987; Finn IM et al. 2007) preserving the preferred direction (Fig. 1E and Suppl. Fig. 2). Second, contrast gain modulations vary widely with contrast across cells (Fig. 2B and Suppl. Fig. 2B) as shown in previous studies (Albrecht DG and DB Hamilton 1982; Ohzawa I *et al.* 1982, 1985; Ledgeway T et al. 2005; Peirce JW 2007; Sani I *et al.* 2013). This is underscored by the fact that ∼28% cells responded stronger to 40% contrast than 100% contrast (*α* > 1) in our data (Fig. 2B, *left panel*). This is commensurate to prior reports (Peirce JW 2007; Sani I *et al.* 2013), including 20-28% such cells identified in monkey V1/2 (Peirce JW 2007) and V4 (Sani I *et al.* 2013).

Our study demonstrated that the population code for grating direction-of-motion deviates significantly from invariance when stimulus contrast is changed. This is manifested i) by the degradation of decoding performance when training/testing decoders from trials taken across contrasts (Fig. 2A) and ii) by the relatively large fraction of distinct cells that contribute to direction-decoding at different contrasts (Fig. 3). Furthermore, controlling for different noise levels and noise correlations across contrasts preserved these conclusions (Fig. 4). Finally, our results were consistent across different stimulus sizes, suggesting they were not a trivial by-product of differences in contrast-dependent spatial integration (Suppl. Fig. 5).

The fact that optimal population codes vary with visual contrast raises the question whether the brain need implement a different code depending on contrast. It is likely that the population code changes continuously with contrast, following a trajectory in a high-dimensional space determined by the cells’ contrast gain response functions. We argue that the heterogeneity of contrast gain response functions across the cell population is the main reason behind the failure of contrast invariance (Fig. 2 and Suppl. Fig. 1). Similar to May and Zhaoping (May KA and L Zhaoping 2011), we hypothesize that the diversity of contrast gain responses across cells may confer an advantage as the same group of neurons is able to encode stimulus orientation/direction and contrast simultaneously. Alternatively, strict contrast invariance would imply that a different group of cells is needed to encode visual contrast itself. To our best knowledge, no cells have been found that are exclusively contrast selective in visual cortex.

### Reconciliation with Prior Results

Prior studies concluded that neuronal populations encode orientation/direction in a contrast-invariant manner. Busse et al. claimed that the population orientation tuning functions (POTF) were contrast invariant (Busse L *et al.* 2009). However, POTFs were calculated by averaging the responses of cell groups with similar orientation preference after normalizing individual cell responses. This method quenches the large diversity of contrast gain responses across cells, which we argue is the source of failure of contrast invariance, and is therefore in no direct conflict with our results. Similarly, Berens et al.(Berens P *et al.* 2012) reported that pooling weights, which were used for linear population decoding, are preserved across contrasts. However, pooling weights are again averaged across cells with similar orientation preference, thereby smoothing out potential heterogeneities.

Another study (Graf AB *et al.* 2011) suggests that neural population direction code is invariant across contrasts even though they report substantial degradation for cross-contrast linear population decoding, a result similar to that shown in our study. To justify this conclusion they assumed that some cells responding vigorously to high contrast do not respond at low contrast, degrading decoding performance at low contrast when the high contrast data were used to train decoders. Although this phenomenon inevitably happens at low enough contrast, it actually does not explain the observations we make here: 1) We observed degradation in cross-contrast performance even when training at low and testing at high contrast, which cannot be readily explained by the above mechanism, 2) we explicitly chose to compare cross-contrast decoding across a smaller contrast transition (100% to 40% and vice versa), thus minimizing the potential problem discussed in (Graf AB *et al.* 2011), 3) we did in fact observe a substantial number of cells that do respond stronger at lower contrast, and therefore contribute more substantially to decoding at lower versus higher contrasts (Figs 1-3), and finally 4) controlling for the signal to noise ratio of responses preserved out conclusions (Fig. 4).

### Spontaneous fluctuations in population activity preserve the population code

Neural responses vary during the repeated presentation of identical stimuli because they are modulated by internal inputs. Such inputs could represent changes of brain state (Niell CM and MP Stryker 2010; Polack PO *et al.* 2013; Fu Y *et al.* 2014; Reimer J *et al.* 2014; McGinley MJ, SV David, *et al.* 2015; Vinck M *et al.* 2015) or simply reflect spontaneous modulations occurring within a stable brain state. Internal inputs that cause modulations reflected in global population activity and correlation structure (Lin IC et al. 2015; Okun M et al. 2015; Rabinowitz NC et al. 2015; Scholvinck ML et al. 2015) may impact population coding (Arandia-Romero I et al. 2016).This raises the question of how population codes “survive” the ever-present modulations driven by internal inputs that may have little to do with the “external” stimulus being encoded.

Our study investigated this question and found that “internal” and “external” inputs have different effect on population codes. Specifically, we found that the population code for direction-of-motion is largely preserved across spontaneous internal input modulations (Figs. 5A-D), but not across changes in stimulus contrast that produce commensurate or smaller firing rate variations (Fig. 5F). We suggest that the reason for this difference is that internal input fluctuations appear to modulate gain more homogeneously across the population of layer 2/3 area V1 cells (Fig. 5E). This is also reflected in the mean signal-to-noise ratio across cells, which is preserved regardless of the different population-activity-levels induced by internal state (Fig. 5G). Further analysis on the most informative cells also revealed that changes in spontaneous population activity preserve population codes (Fig. 6). Cells selected as most informative at a different contrast had much larger gain changes than the ones selected at a different PAL (Suppl. Fig. 8). Our results are in general agreement with two recent studies, which found that: 1) with spontaneous fluctuation of population activity does neuronal activity co-modulate, which does not depend on the similarity of the cells’ preferred orientation (Okun M *et al.* 2015), and 2) brain-state related neuronal fluctuations that occur spontaneously and are thereby uncorrelated with the stimulus do not impact decoding performance (Moreno-Bote R et al. 2014). Given that the cells’ contribution to direction decoding was highly correlated across different PALs (Suppl. Fig. 7), these results suggest that population codes may be largely shared across internal states that modulate neuronal responses along multi-dimensional trajectories uncorrelated with the signal change.

Overall, a spontaneous change in population activity level (i.e., from the upper to the lower 50^th^ percentile) affects the population code much less than a change in stimulus contrast from 100% to 40% (see Figs. 5-6). This occurs even though L2/3 aggregate firing rates change more in the former case. We stress that we do not necessarily imply that internal modulations leave population codes perfectly invariant. In fact, we report that population codes between high and low PALs show a small but significant difference (Suppl. Fig. 7). Furthermore, we cannot exclude the possibility that there may be a small sub-population of cells that behave differently under internal fluctuations, as argued recently by Arandia-Romero et al. (Arandia-Romero I *et al.* 2016). However, when considering all neurons differently behaving with population activity, the neural code is robust to spontaneous modulation in population activity (Fig. 6).

We note that population activity level, though it is known to reflect certain behavioral and brain-state changes (Niell CM and MP Stryker 2010; Lee SH and Y Dan 2012; Polack PO *et al.* 2013; Fu Y *et al.* 2014; Reimer J *et al.* 2014), may have limitations as a surrogate measure of the internal input state. However, we believe that it is sufficient to support our claims. We base this on prior results showing that spontaneous neuronal population activity levels (PAL) co-modulate strongly with simultaneously recorded neighboring neuropil activity as well as EEG and EcoG activity (Kerr JN et al. 2005; Lee S *et al.* 2017), often used to assess brain state (Lee SH and Y Dan 2012). It is also important to note that we investigated the effect of internal input fluctuations occurring *<u>spontaneously</u>*, while animals are sedated or in the quiet-wakefulness state. It is interesting to consider in the future how active changes in brain state, such as modulations of attention, impact the conclusions we have drawn here.

## Conclusion

The present study demonstrates how two different neuronal gain modulation mechanisms, one “external” and the other internal, influence population coding. Gain responses induced by changes in stimulus-contrast as an example of external input are heterogeneous across cells and reshape population codes, whereas gain responses induced via the fluctuation of internal inputs are more homogeneous and do not. The heterogeneous gain modulation works particularly for contrast-specific population codes by increasing response gains for informative cells selected at a contrast, but not at another contrast. This observation has implications for visual information encoding, and argues that the brain strives to maintain stability of neural encoding in the face of markedly fluctuating internal inputs.

## Acknowledgments

This work was supported by a Simons Foundation Pilot Award and R21 NS088457 from NINDS to SS. The authors thank Drs Emmanouil Froudarakis, Jake Reimer for helpful input regarding awake experiments, and Dr. Ryan Ash for editing suggestions. The authors declare that no competing interests exist. S.L and S.M.S conceived the project. J.P. prepared animals and performed surgery with optimizing virus injection and chronic window. S.L. built the experimental setup, performed experiments, analyzed the data, and wrote the original draft with input from J.P. S.L and S.M.S edited the draft.

